# D-serine- and *p*-fluorophenylalanine-resistant mutants of *Physcomitrium patens* are defective in amino acid uptake

**DOI:** 10.1101/2023.08.11.552859

**Authors:** Stephen Albone, David J Cove, Neil W Ashton

## Abstract

Two amino acid analogue-resistant lines of *Physcomitrium* (formerly *Physcomitrella*) *patens, DSR8* and *PFR4*, are resistant to a range of D-amino acids and an inhibitory concentration of L-lysine. Both are defective in the uptake of [^35^S]-L-methionine. Uptake by the wild-type line is pH-dependent (decreasing with raised external proton concentrations) and is depressed by dinitrophenol and by ammonium ions. We discuss the possible involvement of an active proton-general amino acid antiport pump in the plasma membrane.

## Introduction

Previously we have shown that the growth of wild-type *Physcomitrium* is inhibited by the inclusion of either 10 mM D-serine or 550 μM p-fluorophenylalanine (pfpa) in the culture medium, producing small protonemal colonies consisting predominantly of chloronemata. We have also reported the isolation, following mutagenesis with *N*-methyl-*N′*-nitro-*N*-nitrosoguanidine or ethyl methane sulphonate, of D-serine- and pfpa-resistant mutants. The degree of resistance varied but some lines grew equally well in the presence or absence of the amino acid analogue. The most resistant lines were resistant to both analogues (Ashton and Cove 1977). Here we report the responses of two of the most vigorous of the amino acid analogue-resistant lines, *DSR8* and *PFR4*, to a range of D-amino acids and an inhibitory concentration of L-lysine. We also examined the ability of these mutants to accumulate [^35^S]-L-methionine.

## Materials and Methods

### Plant material and culture conditions

For growth tests, plants were grown axenically on solid ABC medium (Knight et al. 1988) at approx. 22-25 °C in continuous white light for approximately 4 weeks. Amino acids and amino acid analogues were added as solids to the medium prior to autoclaving. Agar media containing 5mM L-aspartate, D-glutamate or L-cysteine did not solidify and were discarded. Therefore, 1 mM was the highest concentration at which these amino acids were tested. Protonemal tissue used for testing the ability of wild-type and mutant lines to take up [^35^S]-L-methionine was grown on solid medium with or without di-ammonium (+) tartrate (5 mM) overlaid with sterile, porous cellophane discs (Grimsley et al. 1977), and inoculated with fragmented moss.

### Uptake assay

[^35^S]-L-methionine, with an original specific activity of approximately 1μCi.μmol^-1^, was added to uptake incubation medium to give a starting radioactivity of approximately 10,000 -70,000 cpm per 20 μl sample depending on the particular experiment. 1-4 g of approximately one to two week old moss protonemal tissue depending on the particular experiment, which had been pre-soaked in buffer and filtered, was added to each incubation vial containing 5 ml of incubation medium. Incubation medium was buffered with 50 mM 2-(N-morpholino)ethanesulfonic acid (MES) at pH 6 or 7 or 50 mM citrate-KOH at pH 4 or 5. 100nM or 250 nM cold carrier L-methionine was included in the incubation medium for the pH experiment and the experiment with *DSR8*, respectively In some cases, 1 mM dinitrophenol (DNP) or 5 mM di-ammonium (+) tartrate was included in the buffer solution. Uptake experiments were performed at room temperature in ambient light. Following incubation of between 0.25 and 5 h, 20 μl samples of incubation medium were added to 10 ml of Optiphase X scintillation fluid and remaining radioactivity measured by 3 or 5 minute counts in a TRI-CARB 4430 liquid scintillation counter. Uptake by the moss tissue was calculated as the percentage of cpm removed from the incubation medium (((initial cpm – remaining cpm)/initial cpm) X 100). Calculations and graphs were made using Microsoft Excel. Data were fitted to logarithmic curves.

## Results

### Growth tests with amino acid analogues and an inhibitory concentration of L-lysine

The wild-type line is strongly inhibited by 500 μM and 5 mM D-serine (Fig. 1B), 500 μM and 1 mM pfpa (Fig. 1C), 5mM D-phenylalanine (Fig. 1D), 500 μM and 5 mM D-alanine (Fig. 1E), 2mM and 5 mM D-leucine (Fig. 1F), 1mM and 5 mM D-methionine (Fig. 1G) and 5 mM L-lysine (Fig. 1H). It is slightly sensitive to 500 μM D-glutamine (Fig. 1I) and insensitive to 1mM and 5 mM D-valine, 1mM and 5 mM L-arginine, and 1 mM D-glutamate.

**Figure 1.**
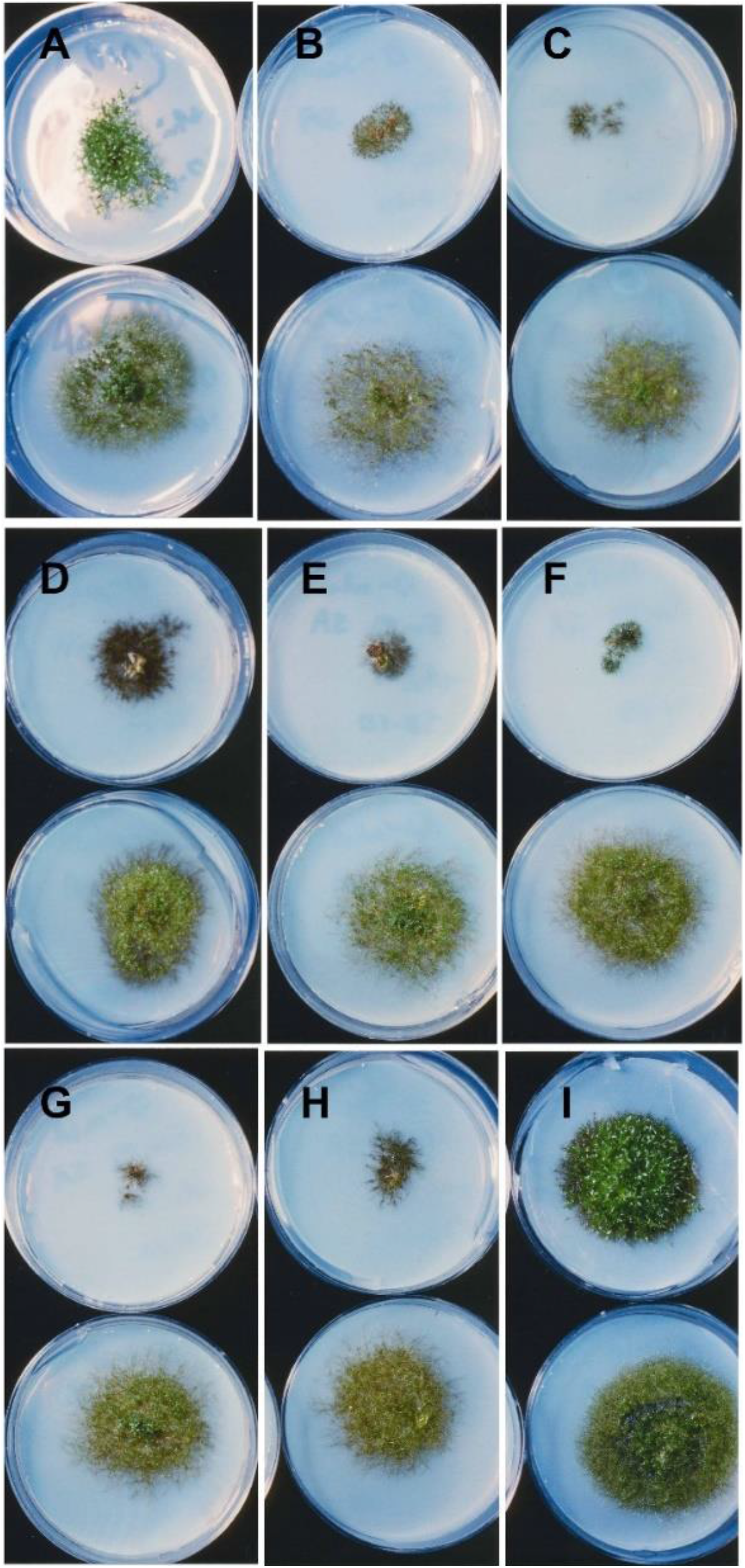
Growth responses of wild-type and *DSR8* to D-amino acids, *p*-fluorophenylalanine and an inhibitory concentration of L-lysine. Wild-type (upper rows) and *DSR8* (lower rows) grown on minimal medium **(A)** and medium containing 5 mM D-serine **(B)**, 500 μM *p*-fluorophenylalanine **(C)**, 5 mM D-phenylalanine **(D)**, 5 mM D-alanine **(E)**, 5 mM D-leucine **(F)**, 5 mM D-methionine **(G)**, 5 mM L-lysine **(H)** and 500 μM D-glutamine **(I)**. Petri plates measure 6 cm in diameter.

*DSR8* and *PFR4* are strongly resistant to 500 μM and 5 mM D-serine (Fig. 1B), 500 μM and 1 mM pfpa (Fig. 1C), 5mM D-phenylalanine (Fig. 1D), 500 μM and 5 mM D-alanine (Fig. 1E), 2mM and 5 mM D-leucine (Fig. 1F), 1mM and 5 mM D-methionine (Fig. 1G) and 5 mM L-lysine (Fig. 1H). *DSR8* and wild-type grown on 500 μM D-glutamine (Fig. 1I) have a similar appearance but *DSR8* is slightly larger and greener than wild-type, which has darkly pigmented peripheral protonemata. The mutant lines and wild-type respond similarly on 1mM and 5 mM D-valine, 1mM and 5 mM L-arginine and 1 mM D-glutamate. Therefore, we cannot reach any conclusions about the resistance of *DSR8* and *PFR4* to these latter compounds.

### [35S]-L-methionine uptake by wild-type, *DSR8* and *PFR4*

The uptake of [^35^S]-L-methionine by the wild-type line is pH-dependent (decreasing with raised external proton concentrations), being greater at pH 6 and 7 than at 4 and 5 (Fig. 2A). It is depressed by dinitrophenol (Fig. 2B) and by ammonium ions present in the culture medium (Fig. 2C) or the uptake incubation medium (Fig. 2D). Both *DSR8* (Fig. 2B) and *PFR4* (Fig. 2C) are defective in the uptake of [^35^S]-L-methionine.

**Figure 2.**
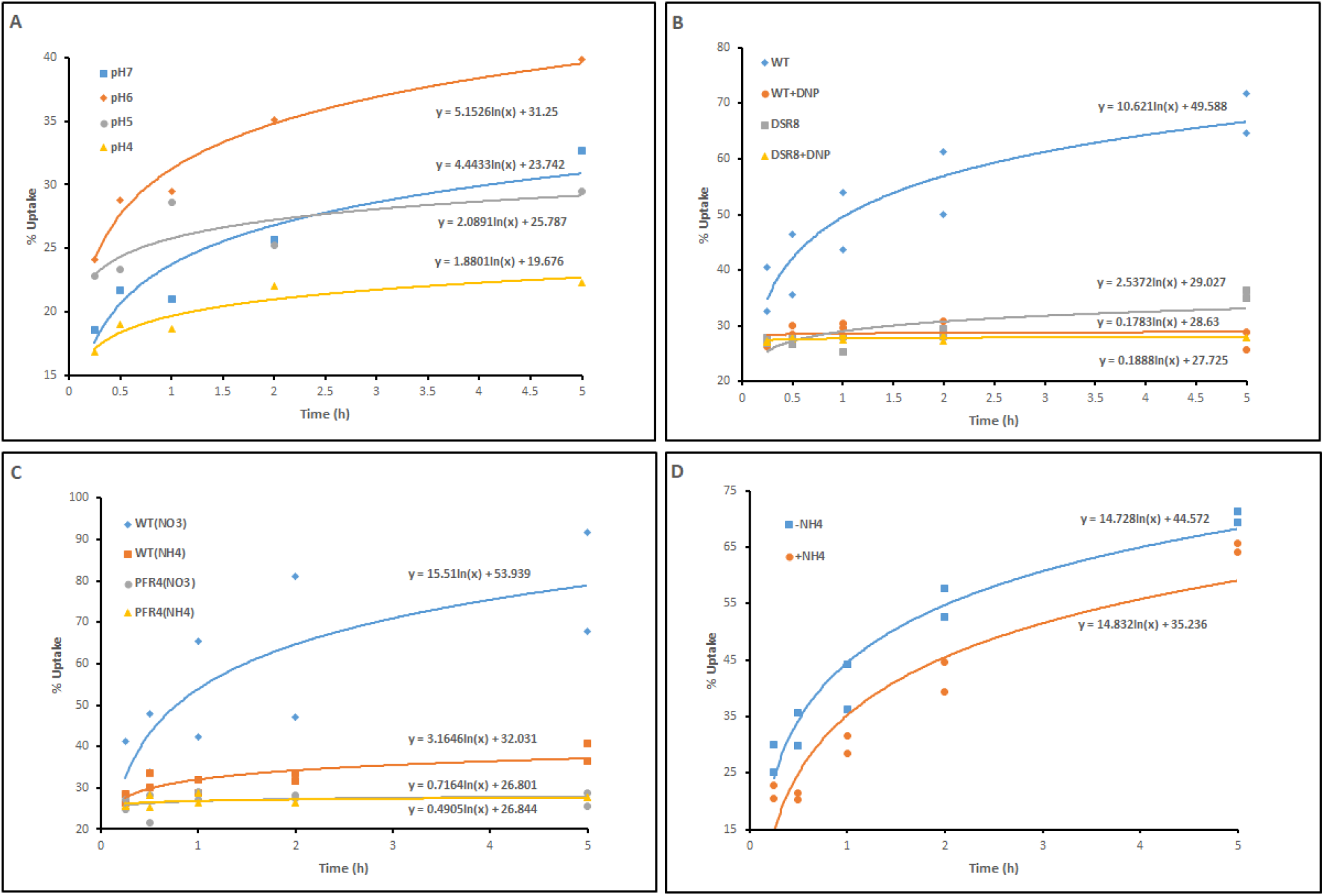
Uptake of [^35^S]-L-methionine by wild-type, *DSR8* and *PFR4*. (A) Wild-type incubated in 5 mM MES buffer (pH 6 and 7) or 5 mM citrate-KOH buffer (pH 4 and 5). (B)Wild-type and *DSR8* incubated in 5 mM MES buffer (pH 6) with and without 1 mM dinitrophenol. (C) Wild-type and *PFR4*, grown on medium with nitrate as sole nitrogen source or with additional ammonium, incubated in 5 mM MES buffer (pH 6). (D) Wild-type, grown on medium with nitrate as sole nitrogen source, incubated in 5 mM MES buffer (pH 6) with and without 5 mM di-ammonium (+) tartrate. Functions defining the fitted logarithmic trendlines are displayed above or below the lines.

## Discussion

The pH dependence of [^35^S]-L-methionine uptake (decreasing with raised external proton concentrations) and inhibition by DNP implies the presence of an amino acid-proton antiport carrier in the plasma membrane (PM) that requires a source of energy. This finding was unexpected since most amino acid transporters in plants appear to be amino acid-proton symports (Rensch et al. 2007, Tegeder and Rensch 2010, Yao et al. 2020 and references therein). In *Riccia fluitans*, for example, increasing the external concentration of protons between pH 9 and 4.5 stimulates the uptake of basic amino acids and glutamate uptake occurs only at acidic pHs. More in keeping with our findings, histidine uptake has a pH optimum between 5.5 and 6 (Johannes and Felle 1985). Additional studies using other labelled amino acids will be needed to clarify this situation.

While we do not know the basis of the observed inhibition of amino acid uptake by ammonium, it is not surprising since ammonium is the best source of inorganic nitrogen for plants and is converted in one step to glutamate, which is a starting point for the synthesis of other amino acids and nitrogen-containing compounds.

During the last two decades, it has been discovered that plants possess a large number of different amino acid transporters with specialised functions within the plant including uptake of amino acids by root epidermal cells and root hairs from the rhizosphere and loading of xylem and phloem for transport to amino acid sinks including leaves, flowers, seeds and root tips (Tegeder and Rensch 2010, Yao et al. 2020). For example, a search of phytozome version 13 reveals that *Arabidopsis thaliana* has 63 amino acid transporters, *Oryza sativa* has 99, *Selaginella moellendorffi* has 79 and *Physcomitrium* has 69. The transporters belong to two main families: Amino Acid/Auxin Permease (AAAP), also known as Amino acid Transporter (ATP), and Amino Acid/Polyamine/organoCation (APC). Each of these has been divided into several sub-families. Thus, The AAAP family comprises general Amino Acid Permeases (AAP), Lysine and Histidine Transporters (LHT), γ-Aminobutyric acid Transporters (GAT), Proline Transporters (ProT), Aromatic and Neutral amino acid Transporters (ANT) and Auxin transporters (AUX). The APC family comprises Cationic Amino acid Transporters (CAT), amino acid/choline transporters and Polyamine-proton Symports (PHS). Names of these transporters is based on their originally assigned selectivity or selectivity in other kingdoms. Therefore, they do not always reflect complete or correct substrate specificity of a specific transporter (Rensch et al. 2007). For example, LHTs were originally described as lysine and histidine selective transporters (Chen and Bush 1997) but were shown later to preferentially transfer neutral and acidic amino acids across the PM with high affinity (Lee and Tegeder 2004, Hirner et al. 2006). An important characteristic of many of these transporters is that they transport a broad range of amino acids. This is especially true for LHTs discussed above and for AAPs, which transport neutral amino acids and glutamate. It is noteworthy that, among *Physcomitrium*’s 69 amino acid transporters, there are 23 AAPs and 4 LHTs. While we do not know specifically which transporter is defective in *DSR8* and *PFR4*, the broad spectrum of amino acids transported by AAPs and LHTs accords well with the mutant lines’ resistance profiles. Although it is simplest to postulate that each of the mutant lines is defective in a single transporter, we can’t exclude the possibility that they possess mutations in two or several genes encoding different amino acid transporters. Another possibility is that they carry a mutation in a regulatory gene controlling the expression of several to many transporters.

### Notes on contributors

Stephen Albone studied as an undergraduate under Professor David Cove at the University of Leeds, UK. He subsequently pursued a career in educational research and is presently engaged as a technical consultant in the evaluation of humane education programmes.

David Cove is Professor Emeritus of Genetics at the University of Leeds, UK. As a graduate student in 1960, he used the fungus *Aspergillus nidulans* for biochemical genetic studies. He began to develop *Physcomitrella* (now *Physcomitrium*) *patens* as a model system for developmental genetic studies in the 1970s. Neil Ashton was his first graduate student to use the moss.

Neil Ashton is Professor Emeritus of the University of Regina. He began his research on *Physcomitrella patens* in 1980 as a graduate student under the supervision of Dr. David Cove in the Genetics Department, University of Cambridge and has continued studying this model plant system to the present day.

